# A glucose-stimulated BOLD fMRI study of hypothalamic dysfunction in mice fed a high-fat and high-sucrose diet

**DOI:** 10.1101/2020.03.21.001149

**Authors:** Adélaïde A. Mohr, Alba M. Garcia-Serrano, João P.P. Vieira, Cecilia Skoug, Henrik Davidsson, João M.N. Duarte

**Affiliations:** Department of Experimental Medical Science, Faculty of Medicine, Lund University, Lund, Sweden; Wallenberg Centre for Molecular Medicine, Lund University, Lund, Sweden

**Keywords:** glucose sensing, neuronal stimulation, functional imaging, diet-induced obesity, hypothalamus

## Abstract

The hypothalamus is the central regulator of energy homeostasis. Hypothalamic neuronal circuits are disrupted upon overfeeding, and play a role in the development of metabolic disorders. While mouse models have been extensively employed for understanding mechanisms of hypothalamic dysfunction, functional magnetic resonance imaging (fMRI) on hypothalamic nuclei has been challenging. We implemented a robust glucose-induced fMRI paradigm that allows to repeatedly investigate hypothalamic responses to glucose. This approach was used to test the hypothesis that hypothalamic nuclei functioning is impaired in mice exposed to a high-fat and high-sucrose diet (HFHSD) for 7 days. The blood oxygen level-dependent (BOLD) fMRI signal was measured from brains of mice under light isoflurane anaesthesia, during which a 2.6 g/kg glucose load was administered. The mouse hypothalamus responded to glucose but not saline administration with a biphasic BOLD fMRI signal reduction. Relative to controls, HFHSD-fed mice showed attenuated or blunted responses in arcuate nucleus, lateral hypothalamus, ventromedial nucleus and dorsomedial nucleus, but not in paraventricular nucleus. In sum, we have developed an fMRI paradigm that is able to determine dysfunction of glucose-sensing neuronal circuits within the mouse hypothalamus in a non-invasive manner.

## Introduction

Brain function requires continuous glucose supply (Sonnay *et al*., 2017), which is ensured by a glucose sensing network across several brain regions (Pozo & Claret, 2018). The hypothalamus is pivotal in the central glucose sensing, in addition to controlling other basic life functions such as feeding behaviour, thermoregulation, sleep or fear response (Pozo & Claret, 2018; Timper & Brüning, 2017). Different hypothalamic nuclei are involved in regulating peripheral metabolism to maintain glucose homeostasis (figure 1), namely the arcuate nucleus (ARC), the lateral hypothalamus (LH), the ventromedial nucleus (VMN), the dorsomedial nucleus (DMN) and the paraventricular nucleus (PVN) (Yeo & Heisler, 2012). The ARC and VMN, as well as brainstem and corticolimbic structures, contain glucose-sensing neurons (Pozo & Claret, 2018): high glucose levels depolarize glucose-excited neurons via glucokinase (GK) and ATP-mediated closure of ATP-sensitive K^+^ channels, and low glucose concentration depolarises glucose-inhibited neurons namely via AMPK-dependent closure of Cl^-^ channels.

**Figure 1.**
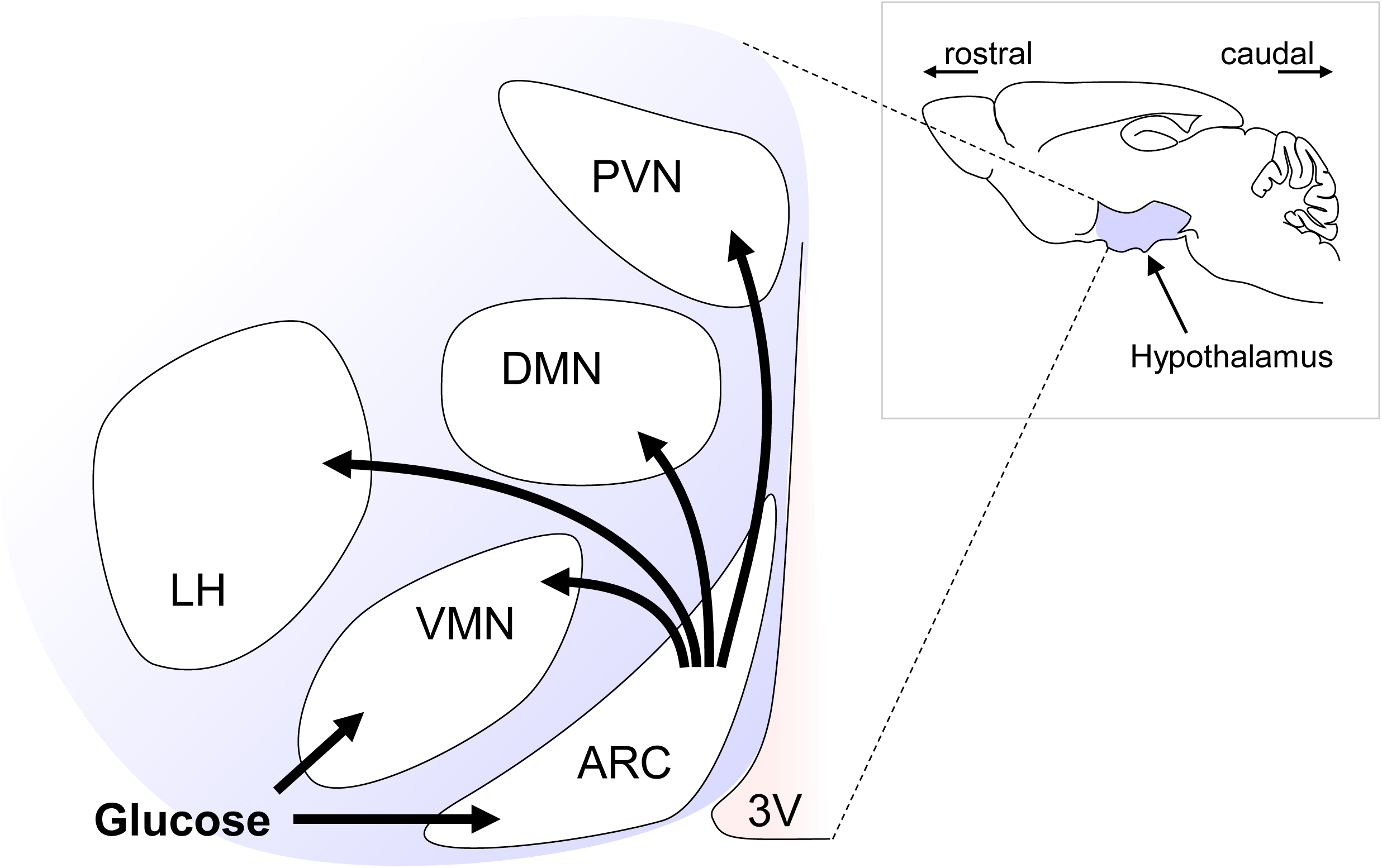
Schematic representation of hypothalamic nuclei activation in response to glucose. Abbreviations: PVN, paraventricular nucleus; DMN, dorsomedial nucleus; ARC, arcuate nucleus; LH, lateral hypothalamus; VMN, ventromedial nucleus; 3V, third ventricle.

Key to the control of appetite, energy expenditure, and glucose homeostasis is the hormonal regulation of the melanocortin system, which consists of two functionally antagonistic neuronal populations in the ARC (Pozo & Claret, 2018; Timper & Brüning, 2017): one subset of neurons expresses the orexigenic neuropeptides (appetite-stimulating) agouti-related peptide (AgRP) and neuropeptide Y (NPY), the second subset expresses the anorexigenic peptides (appetite-suppressing) pro-opiomelanocortin (POMC) and cocaine and amphetamine regulated transcript (CART). These signals are then integrated by secondary neurons in other hypothalamic nuclei, namely the PVN, VMN, DMN and LH, as well as in extra-hypothalamic areas (Timper & Brüning, 2017). ARC neurons also receive feedback from other hypothalamic nuclei, resulting in a finely tuned response to control food intake (Waterson & Horvath, 2015).

Hypothalamic function has been measured non-invasively by magnetic resonance imaging (MRI). Smeets and colleagues reported a prolonged dose-dependent decrease of the functional MRI (fMRI) signal in the hypothalamus after a glucose load (Smeets *et al*., 2005). This response started shortly after subjects drank the glucose solution and lasted for at least 30 min, and was most pronounced in the upper anterior hypothalamus. In the upper posterior hypothalamus, the signal decrease was independent of the glucose load, and no effect was found in the lower hypothalamus. Such hypothalamic responses have also been reported elsewhere (van Opstal *et al*., 2005). Conducting such studies in animal models has been challenging due to breathing or uncontrolled animal movement that generates image artifacts, or biased by the employment of anaesthesia. Nevertheless, Mahankali and colleagues found a negative BOLD fMRI response elicited by i.p. glucose administration the hypothalamus of rats under α-chloralose anaesthesia (Mahankali *et al*., 2000). Lizarbe and colleagues have also previously observed T_2_*-weighted contrast changes in the hypothalamus of mice in an image acquired during glucose infusion *versus* baseline (Lizarbe, Fernández-Pérez *et al*., 2019). While this study lacked temporal resolution to evaluate dynamic responses of hypothalamic nuclei, we aimed at establishing an experimental setup for evaluating the function of the hypothalamus in real time using glucose-induced fMRI.

Metabolic syndrome and obesity are associated to hypothalamic dysfunction through mechanisms that include neuroinflammation and the subsequent insulin and leptin resistance of neurons, which disrupts sensing of metabolic cues, and further promotes food intake and body weight gain (Timper & Brüning, 2017). Hypothalamic inflammation is actually an early event in the development of the metabolic syndrome upon overfeeding, and mice exposed to a high-fat diet show an inflammatory response within one day, which is proposed to play an important role on the subsequent hypothalamic neurodegenerative process (*e.g.* Thaler *et al*., 2012). Therefore, we further investigated the impact of short-term high-fat and high-sucrose diet (HFHSD) exposure on the fMRI-detected hypothalamic response to glucose.

In sum, the goal of this work is 2-fold: (*i*) to develop a paradigm for non-invasively assessing hypothalamic functioning in mice; (*ii*) to map hypothalamic dysfunction upon short-term high-fat and high-sucrose feeding.

## Material and Methods

### Animals

All procedures on animals were approved by the Malmö/Lund Committee for Animal Experiment Ethics (permit number 994/2018), and are reported following the ARRIVE guidelines (Animal Research: Reporting *In Vivo* Experiments, NC3Rs initiative, UK). Male C57BL/6J mice were obtained from Taconic (Ry, Denmark) at 8 weeks of age and allowed to acclimatise to the animal facility for one week. Mice were housed in groups of 4-5 on a 12-hour light-dark cycle with lights on at 07:00, room temperature at 21-23 °C, humidity at 55-60%, and access to tap water and food *ad libitum*. Controls were fed a low-fat diet with 10% kcal from fat, 20% kcal from protein and 70% kcal from carbohydrates (D12450J, Research Diets, New Brunswick, NJ, USA). HFHSD-exposed mice were fed a high-fat diet with 60% kcal from fat, 20% kcal from protein and 20% kcal from carbohydrates (D12492, Research Diets), and were also supplemented with a 20% (w/v) sucrose solution in addition to tap water. The sample size for this study was estimated with the resource equation method (Festing & Altman, 2002).

### Glucose tolerance test (GTT)

Animals were fasted for 6-8 hours starting at 7:00, and then received a load of glucose i.p. (2 g/kg; prepared as 20% (w/v) solution in sterile saline, BRAUN, Melsungen, Germany). Glycaemia was monitored with a glucometer (Accu-check Aviva, Roche, Stockholm, Sweden) from 1-µL tail tip blood samples before the glucose load, and afterwards at 15, 30, 60, 90 and 120 minutes (Soares *et al*., 2018). A 20-µL blood sample was collected from the saphenous vein for fasting insulin quantification (ELISA kit #10-1247-01 from Mercodia, Uppsala, Sweden).

### MRI experiments

On the day of scanning, access to food was terminated at 8:00, and experiments took place from 14:00 to 16:00.

Mice were anesthetized with isoflurane by induction at 3.5% and maintenance at 2% vaporized in a 1:2 O_2_:N_2_O gas mixture. A PE50 line (Warner Instruments, Hamden, CT-USA) was inserted in the mouse trachea for mechanical ventilation, and a cannula was placed i.p for glucose or vehicle infusion. Then mice were positioned on the MR-compatible bed with a teeth bar and two ear bars for stereotaxic head fixation. The mouse was then fitted into a home-build mask that connected the tracheal line to the MR-compatible mechanical ventilator (MRI-1, CWE, Ardmore, PA, USA). When breathing was assisted by mechanical ventilation, mice received an i.p. injection of 0.1g/kg of pancuronium bromide (Sigma-Aldrich, Schnelldorf, Germany) diluted in saline. The ventilation rate was kept at 90 breaths per minute with a volume of 1.9-2.2 mL and 50% inspiration rate. Isoflurane level was then decreased to 0.7% (Sonnay *et al*., 2018), and the mice were then inserted into the MRI scanner. Body temperature was continuously monitored with a rectal temperature probe (SA Instruments, Stony Brook, NY, USA) and maintained at 36-37 °C with a warm water circulation system. Breathing rate was also confirmed by the SA Instruments monitoring system.

MRI experiments were performed on a Bruker BioSpec 9.4T equipped with a ^1^H quadrature transmit/receive MRI cryoprobe (Baltes *et al*., 2009). A localizer MRI sequence was first used to confirm good positioning of the animal in the scanner and allow for eventual adjustments. Then a set of T_2_-weighted images over the whole brain was acquired for anatomical reference using Rapid Imaging with Refocused Echoes (RARE) sequence with repetition time (TR) of 3.5 s, echo time (TE) of 33 ms, 0.5 mm slice thickness, image size of 320×320 and field of view (FOV) of 14×14 mm^2^. These images were used to position the slices for fMRI. BOLD fMRI data were acquired during 20 minutes using gradient echo-echo planar imaging (GE-EPI) sequence with TR=2 s and TE=18 ms (Sonnay *et al*., 2016). Signal was acquired from four 0.5-mm slices encompassing the whole hypothalamus, resolution of 0.104×0.109 mm^2^ and FOV of 14×10 mm^2^. After 5 minutes of fMRI scanning, either 2.6 g/kg of glucose (from a 30% solution in saline) or equivalent saline volume was injected i.p. during 3 minutes. Saline infusion was usually performed in the same mouse before or after the glucose infusion. The order of the two infusions was randomly chosen to eliminate systematic errors, and there was an interval of at least 30 min between the two infusions. After the experiment, mechanical ventilation and isoflurane anaesthesia were stopped when spontaneous breeding reflexes were observed, and mice returned to their cages.

Mock experiments out of the scanner were conducted in 2 mice to determine the typical glycaemic profile during the glucose-induced fMRI paradigm. Glucose was determined from tail-tip blood samples.

### fMRI data analysis

EPI images were reconstructed and corrected for motion using SPM12 (Statistical Parametric Mapping, London, UK), and the average signal from predefined regions of interest (ROI) in the hypothalamus were extracted (Sonnay *et al*., 2016). Namely, the hypothalamus was separated in four different ROIs: the dorsomedial nucleus (DMN), the paraventricular nucleus (PVN), the arcuate nucleus (ARC) and finally the lateral hypothalamus (LH) plus the ventromedial nucleus (VMN), from which the average signal was extracted. Signals from LH and VMN were analysed as a single ROI due to lack of contrast on anatomical and functional images (LH+VMN). For each experiment, average signal was extracted from the four ROI using the SPM MarsBar toolbox (Brett *et al*., 2002), and normalized to the average signal during the 2 minutes before glucose infusion (baseline signal).

### Immuno-detection of glucose-induced neuronal activation

Mice kept in control diet (n=8) or HFHSD (n=8) for 7 days were fasted as above, anaesthetised with 1.5-2% isoflurane, and then given 2.6 g/kg glucose i.p.. After 20 minutes, half of the animals were sacrificed by decapitation, the brain was rapidly removed and the hypothalamus was dissected, frozen in N_2_ (l), and then extracted for Western blotting as previously (Soares *et al*., 2019). The remaining animals were sacrificed by cardiac perfusion with cold phosphate-buffered saline (PBS; in mmol/L: 137 NaCl, 2.7 KCl, 1.5 KH_2_PO_4_, 8.1 Na_2_HPO_4_, pH 7.4) and then cold 4% para-formaldehyde (PFA) in saline (Duarte et al., 2012).

Western blot analysis was carried out as previously detailed (Lizarbe, Soares *et al*., 2019) using antibodies against cAMP response element-binding protein (CREB; clone LB9, produced in mouse, #ab178322, AbCam, Cambridge, UK; RRID:AB_2827810), phospho-CREB (Ser133) conjugated with AlexaFluor488 (produced in rabbit, #06-519-AF488, Milipore, Temecula, CA, USA; RRID:AB_310153), c-fos (produced in rabbit, #SAB2100833, Sigma-Aldrich; RRID:AB_10600287), and β-actin conjugated with horseradish peroxidase (produced in mouse, #A3854, Sigma-Aldrich; RRID:AB_262011). Secondary antibodies were conjugated with horseradish peroxidase: goat anti-rabbit IgG (#ab6721, AbCam; RRID:AB_955447); goat anti-mouse IgG, #ab6789, AbCam; RRID:AB_955439).

Brains for microscopy were processed as previously described (Duarte *et al*., 2012). Briefly, brains were fixed in phosphate-buffered formaldehyde (Histolab, Askim, Sweden), and then stored at 4 °C in a 30% sucrose solution in PBS. Immunostaining was carried out on 20-µm cryostat-sectioned coronal brain slices. Slices were incubated for 1 hour at room temperature with blocking buffer (PBS containing 5% normal goat serum, 1% BSA and 0.3% Triton X-100), and then overnight at 4 °C with the AlexaFluor488-conjugated antibody against phosphor-CREB (1:50 in blocking buffer). After washing in PBS, slices were mounted with ProLong Glass Antifade (Invitrogen), and examined under a Nikon A1RHD confocal microscope with a 20x Plan Apo objective, NA 0.75 (Nikon Instruments, Tokyo, Japan). Images were acquired with NIS-elements, version: 5.20.01 (Laboratory Imaging, Nikon). Images were processed for nuclei counting in ImageJ (NIH, Bethesda, MD, USA).

### Statistics

Statistical analyses were performed using Prism 8.0.1 (GraphPad, San Diego, California, USA). Results are shown as mean±SEM unless otherwise stated. Data was assumed to follow normality, linearity and equality of variance. Comparison of results was carried out with unpaired, 2-tailed Student t-tests, or with ANOVA followed by Fisher’s LSD post-testing. Significant differences were considered for P<0.05.

## Results

### Glucose-induced fMRI

In pilot experiments, functional MRI experiments with freely breathing mice under 1.5% isoflurane anaesthesia allowed to obtain stable images with limited motion (data not shown). However, we observed unexpected and unpredictable signal variations in bottom areas of the hypothalamus that are closer to the trachea, which were considered to be breathing-associated artifacts. Such artifacts remained during fMRI acquisitions triggered to the breathing rate and, furthermore, variable TR would result in magnetization variations that require additional TR-dependent signal corrections. BOLD fMRI time courses with minimal artifacts were obtained from mice under mechanical ventilation with pancuronium bromide-induced paralysis, as reported by others (Baltes *et al*., 2009). This also allowed us to lower isoflurane anaesthesia level to 0.7%, thus reducing its effects on cerebrovascular reactivity as in previous work (Sonnay *et al*., 2018).

Anatomical images of the rat brain were used to identify the hypothalamus and approximate location hypothalamic nuclei by comparison to a mouse brain atlas (figure 2A). T_2_*-weighted images were obtained from 4 coronal slices encompassing the hypothalamus (figure 2B), and were used to calculate relative BOLD signal variations. These images showed minimal distortions and also provided sufficient contrast to identify the hypothalamus. Upon administration of glucose, the hypothalamus responded promptly with a BOLD signal reduction, which was not observed in other brain areas, such as the cortex (figure 2C). This hypothalamic response appeared to be biphasic, reaching a plateau ∼5 minutes after the beginning of the glucose infusion, and decaying again after ∼10 minutes.

**Figure 2.**
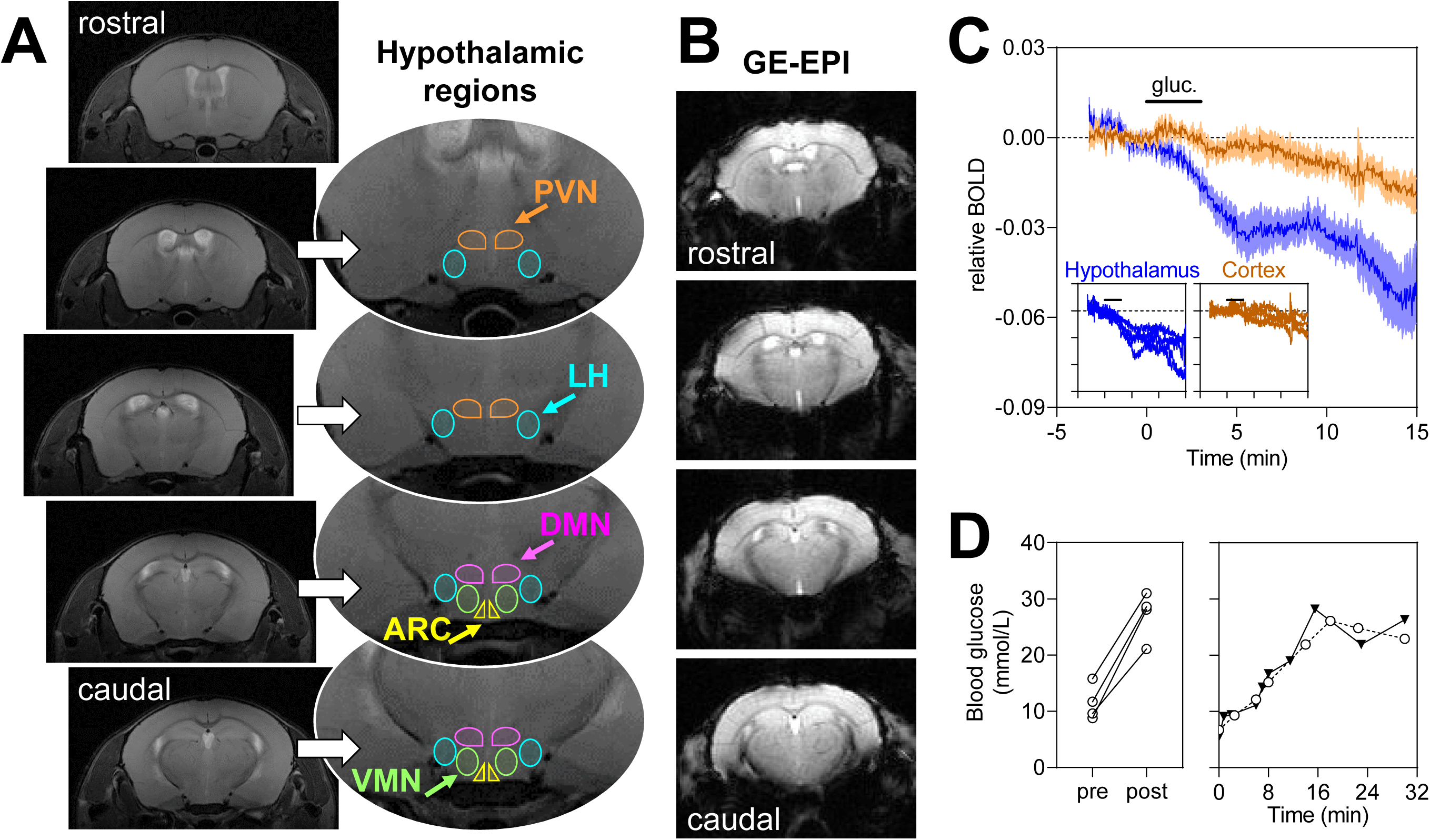
(A) Location of analysed hypothalamic regions overlaid on T_2_-weighted images from a mouse brain: PVN, paraventricular nucleus; DMN, dorsomedial nucleus; ARC, arcuate nucleus; LH, lateral hypothalamus; VMN, ventromedial nucleus. Given the insufficient contrast to distinguish VMN and LH, their voxels were analysed together as LH+VMN. PVN, DMN and ARC were defined as the voxels in the vicinity of the third ventricle. (B) typical T_2_*-weighted contrast in GE-EPI slices encompassing the hypothalamus. (C) Glucose-induced BOLD fMRI response in the hypothalamus of 4 mice is not observable in a ROI of similar size placed in the cortex (shown as mean±SEM). The insets show the trace from each mouse. Glucose infusion for 3 minutes (2.6 g/kg) is indicated by the horizontal bar. (D) Blood glucose levels before (pre) and after (post) the fMRI experiment (left), and in 2 experiments under identical conditions but out of the scanner.

Confirming an effective infusion, blood glucose measured from the tail tip increased from 11.5±1.6 mmol/L before the fMRI experiment to 27.2±2.1 mmol/L at the end of the scan (range of 2-3-fold increase; P=0.002). This increase in blood glucose upon i.p. administration is likely linear during the 15 minutes of experiment, as reproduced in two bench tests (figure 2D).

A continuous BOLD signal drift (∼1% per 15 minutes) was observed across the mouse brain (figure 2C). This was of much lower magnitude than the signal drop after glucose administration, and was also observed in the hypothalamus upon saline infusion (figure 3A). Therefore, it might be that a small fraction of the estimated glucose-induced hypothalamic response includes a glucose-independent signal drift.

**Figure 3.**
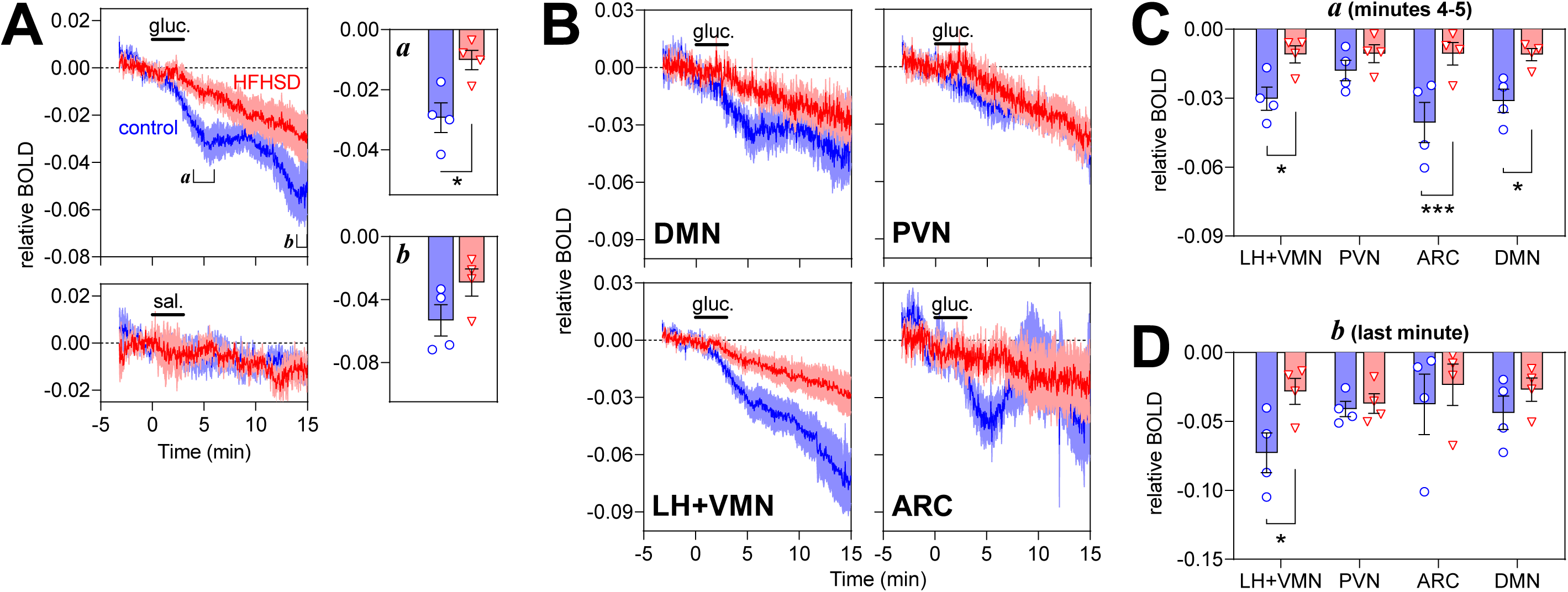
BOLD fMRI signal measured in the hypothalamus (A) or hypothalamic regions (B-D) of control (blue traces/circles) and HFHSD-fed mice (red traces/triangles). (A) BOLD signal changes in response to glucose (top) or saline (bottom). Bar graphs show the BOLD signal averaged during minutes 4 and 5 (*a*) or the last minute of the experiment (*b*). The average BOLD fMRI signal from each area was calculated at 4-5 minutes (C) and minute 15 (D). Glucose (2.6 g/kg) or saline infusion i.p. for 3 minutes is indicated by the horizontal bar. Results are mean±SEM of n=4, and bar graphs show each experimental data-point. * P<0.05, *** P<0.001 in either a Student t-test or Fisher’s LSD post-test performed after significant ANOVA.

### HFHSD effect on hypothalamic responses

Mice exposed to 7 days of HFHSD were of similar weight than controls (27±1 g *versus* 24±1 g; P=0.082), had higher blood glucose (in mmol/L: 9.3±1.0 *versus* 7.0±1.1, P=0.016) plasma insulin (in µg/L: 1.3±0.2 *versus* 0.9±0.1, P=0.039), and similar glucose clearance in a glucose tolerance test (area under the curve in arbitrary units: 10.7±2.1 *versus* 8.1±1.1; P=0.240).

The glucose-induced hypothalamic response was dampened in HFHSD-exposed mice, when compared to controls (figure 3A). That was mainly due to an impairment of the initial response to glucose. When measured at 5 minutes from infusion start, the BOLD signal relative to baseline decreased by 3.0±0.5% in controls and by 1.0±0.3% in HFHSD (P=0.017). After saline infusion, BOLD signal relative to baseline was reduced by only 0.5±0.4% in controls and 0.3±0.2% in HFHSD (P=0.775). When measured at the end of the fMRI experiment (last minute), the BOLD signal relative to baseline decreased by 5.3±1.0% in controls and by 3.0±0.9% in HFHSD (P=0.118). After saline infusion, final BOLD signal was −1.1±0.3% in controls and −1.1±0.5% in HFHSD (P=0.951) relative to baseline.

We further split the BOLD signal originated from different hypothalamic nuclei, namely ARC, LH+VMN, DMN and PVN (figure 2A). All regions responded to glucose infusion but with different magnitudes (figure 3B). The mean relative BOLD slope within the 2 minutes post infusion was −1.3±0.2% per minute in ARC, −0.7±0.1% in LH+VMN, −0.6±0.1% in DMN and −0.3±0.1% in PVN. While this suggests that ARC had the largest initial response (P<0.05 *vs*. other regions in post-tests after significant ANOVA), LH+VMN showed the largest BOLD signal variation from baseline at the end of the experiment (figure 3B). HFHSD exposure had a general effect on glucose-induced BOLD responses, with strongest effects immediately after glucose infusion (ANOVA at 5 minutes after glucose onset: diet F_1,24_=28.27, P<0.001; ROI F_3,24_=1.66, P=0.202; interaction F_3,24_=1.64, P=0.207; figure 3C). When compared to controls, HFHSD mice showed a dampened immediate response to glucose in ARC (P<0.001), LH+VMN (P=0.013) and DMN (P=0.010) but not PVN (P=0.318).

HFHSD exposure had a relatively smaller effect on the BOLD signal change at 15 minutes after the onset of glucose infusion (ANOVA: diet F_1,24_=4.86, P=0.037; ROI F_3,24_=0.89, P=0.460; interaction F_3,24_=0.93, P=0.442; figure 3D). In fact, only the LH+VMN showed a relative BOLD signal difference between HFHSD-fed mice and controls (P=0.021).

The influx of Ca^2+^ into stimulated neurons triggers CREB phosphorylation, which acts as a transcription factor increasing the expression of c-fos mRNA that is then translated into a functional regulator of gene expression (Hudson *et al*., 2018). Levels of pCREB and c-fos were used for confirming neuronal activation (figure 4). Twenty minutes after the glucose load, HFHSD-exposed mice had less pCREB-positive nuclei than controls (repeated measures ANOVA: diet F_1,3_=40.03, P=0.008; ROI F_2,6_=15.09, P=0.005; interaction F_2,6_=0.26, P=0.779; figure 4A). Post-hoc testing identified a HFHSD-induced reduction in the number of pCREB-positive nuclei in all analysed areas (ARC P=0.018, VMN P=0.029, DMN P=0.008). Immunoblotting experiments found no significant change in CREB phosphorylation rate (P=0.067; figure 4B), and a 26±9% reduction of c-fos levels (P=0.029; figure 4C) in the whole hypothalamus of HFHSD-exposed mice, relative to controls.

**Figure 4.**
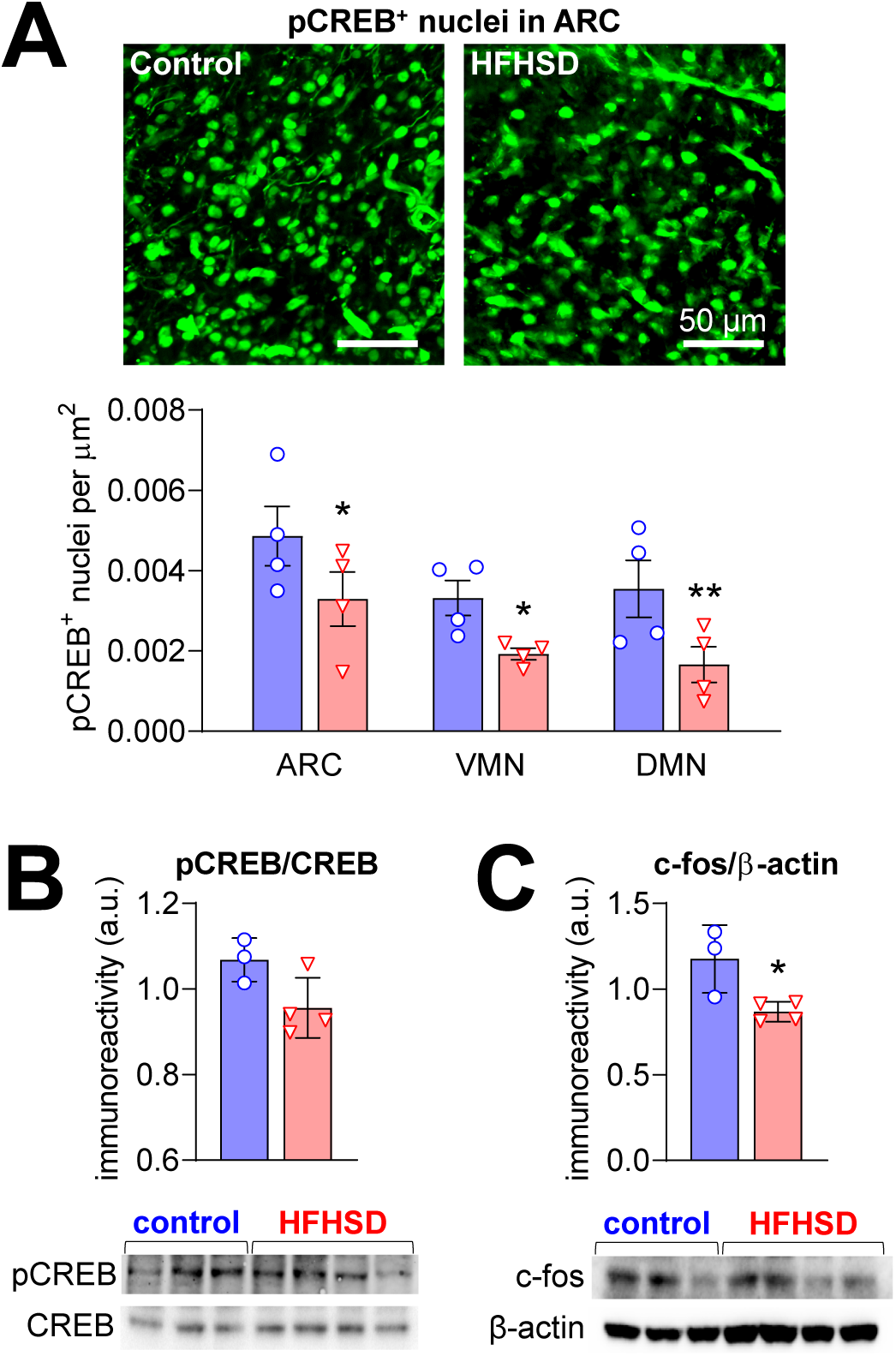
Glucose-induced neuronal activation depicted by CREB phosphorylation and increased levels of c-fos in the hypothalamus. HFHSD-associated reductions in the number of pCREB-positive nuclei were observed in ARC, VMN and DMN (A). Immunoblotting confirmed the reduced CREB phosphorylation (B) and reduced c-fos levels (C) in the hypothalamus of HFHSD-fed mice (red bars/triangles) relative to controls (blue bars/circles). Results are mean±SEM of n=3-4, and are plotted showing each experimental data-point. * P<0.05, ** P<0.01 in either a Student t-test or Fisher’s LSD post-test performed after significant ANOVA.

## Discussion

Hypothalamic neurons sense the nutrient status and integrate signals from peripheral hormones including pancreas-derived insulin and adipocyte-derived leptin to regulate appetite, metabolism and energy expenditure (Pozo & Claret, 2018; Timper & Brüning, 2017). Although understanding the functioning of hypothalamic circuits is key for metabolic diseases research, hypothalamic functional imaging in mouse models remains challenging. In this work we developed an experimental set-up for non-invasively assessing the function of the mouse hypothalamus using fMRI, and found that short-term high-fat and high-sucrose feeding impairs hypothalamic neuronal circuits, particularly in ARC, DMN and VMN+LH.

MRI experiments were performed under light isoflurane anaesthesia, and image artifacts at the level of the hypothalamus could only be prevented by paralysis and mechanical ventilation (Grandjean *et al*., 2017; Sonnay *et al*., 2018). Despite efforts to improve image quality, responses within the ARC remained particularly sensitive to background noise, likely due to its proximity to the trachea. Under these conditions, the hypothalamus responded to glucose with a biphasic BOLD signal reduction – there was a first plateau 5 minutes after glucose infusion followed by a subsequent signal decay. This biphasic response might be caused by different onsets of activation across hypothalamic nuclei and their interactions (figure 1), by signal cancellations due to inhibitory and excitatory responses involving negative and positive BOLD effects (Fukushima *et al*., 2015), or by subsequent hormonal actions, such as insulin (Kullmann *et al*., 2018). We further analysed sub-hypothalamic VOIs in order to dissect their temporal activation. Interestingly, all analysed areas displayed an immediate response upon glucose administration, which was stronger in ARC and weaker in PVN. Since all areas display the typical biphasic response, it is likely that it represents the counter-regulatory hormonal action following the glucose load.

Negative glucose-induced hypothalamic responses have also been observed in humans (Smeets *et al*., 2005; van Opstal *et al*., 2005) and anesthetised rats (Mahankali *et al*., 2000). In mice, others have observed a signal increase in LH and DMN, but not ARC or VMN, in a T_2_*-weighted image acquired during glucose administration relative to baseline (Lizarbe, Fernández-Pérez *et al*., 2019). Interestingly, the authors reported signal reductions in ARC and VMN upon saline administration, which might be attributed to the infusion protocol. One should also note that mice in Lizarbe’s study were exposed to a prolonged fasting period prior experimentation, which is not the case in our work.

To the best of our knowledge, this is the first study investigating the direct response of the hypothalamus after short-term diabetogenic diet feeding in a rodent model. The biphasic responses observed in controls were strikingly attenuated in the hypothalamus of HFHSD-fed mice, especially in the ARC and VMN. This functional impairment was confirmed by reduced molecular markers of neuronal activation (pCREB and c-fos). The ARC is a key element in food intake regulation, since its border with the blood brain barrier is semi-permeable and facilitate passage of blood-stream hormones, and its neurons project to other hypothalamic areas for signal processing and integration (Routh *et al*., 2014; Pozo & Claret, 2018; Timper & Brüning, 2017). The loss of immediate ARC neuronal responses in HFHSD might thus preclude adequate activation the neuronal circuit that allows the hypothalamus to elicit a normal response to glucose.

The measured HFHSD-induced dampening of the BOLD fMRI response to glucose cannot be simply attributed to impaired glucose sensing. The neurovascular coupling that underlies the BOLD signal involves tight metabolic interactions between neurons and glial cells, namely astrocytes (Sonnay *et al*., 2017). The early neuroinflammatory process of the hypothalamus also involves astrogliosis (*e.g.* Thaler *et al*., 2012; Lizarbe, Cherix *et al*., 2019). During this astrogliosis process, the metabolic support from astrocytes to neurons is likely disrupted. For example, our recent work on T2D rats showed altered astrocytic metabolism and impaired glutamine synthesis for glutamatergic neurons (Girault *et al*., 2019; Soares *et al*., 2019), which account for a substantial fraction of hypothalamic cells (Chen *et al*., 2017). Therefore, it is plausible that disrupted neuron-astrocyte metabolic interactions lead to blunted neurovascular coupling, even with preserved glucose sensing at cellular level.

On the other hand, astrocytes are also direct sensors for glucose and fatty acids (Freire-Regatillo *et al*., 2017), leptin (Hsuchou *et al*., 2009) and insulin (Cai *et al*., 2018). Importantly, astrocytic insulin signalling plays a role in glucose-induced neuronal activation (García-Cáceres *et al*., 2016), and obesity impacts astrocyte functions (Hsuchou *et al*., 2009; Cai *et al*., 2018; García-Cáceres *et al*., 2016; Douglass *et al*., 2017). Loss of astrocytic glucose sensing in the hypothalamus may play a role in obesity and T2D development (Freire-Regatillo *et al*., 2017).

## Conclusion

Mouse fMRI experiments are especially challenging in the hypothalamus, which is near an airway that causes major artifacts. While previous functional studies in the mouse hypothalamus lacked temporal resolution to investigate dynamic responses (Lizarbe, Fernández-Pérez *et al*., 2019), we now developed an experimental paradigm for real-time glucose-induced fMRI tasks. This approach is fully non-invasive and can thus be employed for longitudinal mouse studies (Grandjean *et al*., 2017), such as following trajectories of hypothalamic dysfunction.

Although the hypothalamus has a crucial role in regulating food intake, other brain regions also participate in the response to feeding or food cues (Morton *et al*., 2014). Therefore, both resting-state and glucose-stimulated functional connectivity studies can be coupled to the present experimental approach (Grandjean *et al*., 2017).

## Abbreviations

AgRP: agouti-related peptide
AMPK: AMP-activated protein kinase
ARC: arcuate nucleus
BOLD: blood oxygen level-dependent
CART: cocaine and amphetamine regulated transcript
CREB: cAMP response element-binding protein
DMN: dorsomedial nucleus
fMRI: functional magnetic resonance imaging
FOV: field of view
LH: lateral hypothalamus
MC4R: melanocortin 4 receptor
NPY: neuropeptide Y
POMC: pro-opiomelanocortin
PVN: paraventricular nucleus
T2D: type 2 diabetes
TE: echo time
TR: repetition time
VMN: ventromedial nucleus
VOI: volume of interest.

## Acknowledgements

The Knut and Alice Wallenberg foundation, the Medical Faculty at Lund University and Region Skåne are acknowledged for generous financial support. JMND received funding from the Swedish foundation for International Cooperation in Research and Higher Education (BR2019-8508), Swedish Research Council (2019-01130), Diabetesfonden (Dia2019-440), Crafoord Foundation (20190007), and Direktör Albert Påhlssons Foundation. AMGS received funding from the Royal Physiographic Society of Lund, Anna-Lisa Rosenberg Foundation, and Lars Hiertas Minne Foundation. The authors acknowledge the support from the Lund University Diabetes Centre, which is funded by the Swedish Research Council (Strategic Research Area EXODIAB, grant 2009-1039), the Swedish Foundation for Strategic Research (grant IRC15-0067). The authors are grateful to René In’t Zandt and Michael Gottschalk for technical support in MRI experiments. Lund University Bioimaging Centre is gratefully acknowledged for providing experimental resources.

## Author contribution statement

JMND designed the study and wrote the manuscript. All authors performed experiments and analysed data. AAM and AMGS revised and edited the manuscript.

